# LLPSWise - fast and accurate prediction of LLPS constituents

**DOI:** 10.1101/2022.07.25.501404

**Authors:** Mengchen Pu, Biao Fu, Yingsheng J. Zhang

**Affiliations:** StoneWise AI Ltd, Beijing, China

## Abstract

The recent discovery of different types of biological liquid-liquid phase separation (LLPS) systems presents enormous opportunities to uncover underlying biological mechanisms from a single-molecule level to mesoscopic scales and beyond. While progress has been made on the front of computational prediction of LLPS propensity of proteins, the constituent identification of LLPS systems continues to be a pertinent issue that has yet to be addressed. Compiled evidence indicates that the biological mechanism of LLPS involves the employment of different biomolecules into their liquid-liquid separation phase. Since constituent identification relies on experimental approaches, the process is arduous and inefficient. Here, we propose a sequence based LLPS propensity prediction method and scan the human proteome and biochemical pathways to establish a systematic biological view of LLPS. Additionally, we present a fast, accurate method for identifying the constituents of LLPS systems. The source code for our method is available via https://github.com/promethiume/LLPSWise

## Introduction

In the 19th century, cell biologists believed that matters in the cytoplasm were partitioned into liquid-like droplets analogous to suspended chemical mixtures^1^. Present-day, it is observed that P granule in *C. elegans* indeed behaves as antecedently hypothesized^2^. Hitherto, plenty of liquid-like biomolecule condensates were discovered^3-7^. By observation, a certain set of proteins can spontaneously phase separate under both in vitro and in vivo conditions. It has been demonstrated that de novo designed artificial disordered proteins exhibit variable phase behaviors^8^. Phase separation is an intrinsic feature of poly amino acids, particularly for proteins enriched with disordered regions. While the phenomenon of biomolecular LLPS is intriguing, the challenge of relating the phenomenon with biological mechanisms is non-trivial. Alberti et al. have prudently envisaged the possible functional outcomes of LLPS^9^ that are not feasible from membrane-bound organelles. A common belief is that LLPS is limited to macromolecules, however, the partitioning capability of LLPS is well extended to naturally occurring ligands, e.g., the enrichment of ATP and GTP into cGAS condensate^10^ and the selectivity of nuclear condensates exerts on therapeutics^11^. All these observations suggest that LLPS is a meta-mechanism of cellular functions^12,13^, which well complements the functionality of membrane-bound organelles. For fundamental studies or industrial applications, it is highly desirable if one could conveniently know for a given protein, its LLPS propensity; for a given organism, its proteome-wide engagement of LLPS; for certain LLPS system, its constituents. While the first question has been addressed previously^14^, we re-address and integrate it into the constituent identification pipeline. Henceforth, we extend the survey to a human proteome scale and demonstrate our constituent identification method on specific systems.

Identifying specific proteins that could spontaneously phase separate into a condensate/demixed state is a critical and early step in studying LLPS systems. Computational triaging proteins^15,16^ for their LLPS propensities makes it feasible and very efficient for *de novo* identification of the ‘LLPS proteome’. Yet, identifying the LLPS proteome is insufficient for inferring the functionality and biological mechanism of the condensates. Although high LLPS propensity protein is a principal determinant of the LLPS system, a variety of low LLPS propensity proteins must also present to demonstrate the proposed function. Further, some LLPS systems are co-scaffolded by multiple driver proteins^17-19^. We observe that while comparing biological and non-living LLPS systems, the biological system carries a few unique characteristics: it is often specific to its constituents^3,6,7^, intracellular space is crowded with many molecular species which are actively regulated by chemical reactions, and biological LLPS is highly dynamic and an open system that can undergo further phase transition from liquid-like phase to gel-like phase and vice versa^20^.

To relate the LLPS phenomenon with biological mechanisms, the identification of high frequency LLPS system constituents, namely the essential components of physiological LLPS, is a pertinent pursuit in the field. Hitherto it heavily relies on serendipities and experimental techniques. The approach needs to confront arduousness arising from the liquidity and dynamic features of LLPS systems. There were attempts to purify certain LLPS systems, e.g., stress granule (SG) or nucleoli, by seeding cell lysate with an LLPS component and reconstitution^21^ or by chemical cross-linking^22^. Although these are valuable and viable methods, the seeding and reconstitution process does not necessarily reproduce the physiology of LLPS, and it is unclear how the cross-linking process affects the LLPS systems. The interrogation of LLPS constituents otherwise is viable by two low throughput techniques^9,23^. Given the recombinant form of a target protein is available, the in vitro reconstitution method can interrogate its participation into a LLPS system. Furthermore, the contribution of specific sequence regions can be factored by designing different constructs of the target protein. When the target protein is fused with a fluorescent tag, fluorescence imaging is usually the method of choice^24-25^. The throughput of both methods nonetheless cannot catch up with the rapid development in the field. With agreement on the importance of LLPS, an efficient computational method to address the LLPS constituent identification problem is highly desirable.

At first glance, the computational *de novo* identification of LLPS constituents is a formidable task. In this study, we discovered an intriguing mapping relationship between the protein-protein interaction (PPI) network and the LLPS system, and explored it for predicting LLPS constituents, specifically the proteinaceous constituents. We made a few observations before hypothesizing that the PPI network could be mapped to LLPS systems. The observations are as follows: it has long been noticed that intrinsically disordered regions are enriched in eukaryotes and reached a dominant percentage of presence in the human proteome^26^. The functionality of the intrinsically disordered domains (IDDs) is likely to be multifaceted^26,27^, and IDDs are closely related with the phase separation propensity of proteins. The presence of LLPS systems is an early event in the evolution as it is observed in single-celled fungus microorganisms, e.g., *Saccharomyces cerevisiae*. The PPI network is an intrinsic feature of the proteomes, and the presence of LLPS systems is prevalent in both flora and fauna^13,28^. The proteome size of eukaryotes does not necessarily expand from lower to higher organisms, while the percentage of IDDs containing proteins increases almost constantly^26^. LLPS systems and PPI networks have a long history of coevolution, i.e., if a particular protein concomitantly requires both its PPI functionality and LLPS participation to execute its biological function, the two modes of actions need to be compatible with each other and are spontaneously evolved. When an experimental approach is applied for measuring the PPI network from different conditions, the PPI landscape drifts dramatically^29-31^ of which transcriptional activity and expression level of proteins are critical contributing factors. The LLPS mechanism is also powerful in regulating proteins’ spatiotemporal distributions, therefore exerting constraints on the PPI landscape. Conversely, the measured PPIs contain information about this LLPS constraint.

Here, we make accurate predictions on the LLPS network constituents by combining our LLPS propensity prediction and PPI to LLPS mapping algorithm. To triage our LLPS constituent predictions, we compared our results with four relatively well studied LLPS systems: the SPOP-DAXX speckle^32^, a nuclear condensate for proteasomal labeling to the ubiquitination degradation pathway; the cGAS-STING system^33^, a widely known LLPS system which present upon viral DNA or chromosome injuries induction^34^; the SARS-CoV-2 nucleocapsid protein N condensate^35,36^, which carries multiple roles in the viral cycle; and the stress granule (SG)^37^, a membrane-less assemblies of RNA and proteins. Some of these LLPS systems have plenty of identified constituents, however, we are not clear about their exact functionalities. Likewise, we are not providing an exhaustive list of the LLPS constituents, and our prediction should be interpreted as prediction on persistent and core constituents. To our knowledge, this method that can readily facilitate the studies on LLPS systems is the first of its kind.

## Results

### LLPS propensity prediction method benchmark

To evaluate the performance of our predictor based on the statistical indexes of accuracy, F1-score, sensitivity and specificity, we ran a 10-fold cross-validation. Our model achieved the cross-validation training accuracy of 96% ± 1% compared with 89% ± 2% in DeePhase. The statistical index values of the test are shown in Table 1. We also tested our model on the rationally designed A-IDP data set (Supplementary Table 1) and compared our LLPS propensity with the concentration of phase separation transition temperature at 37 °C. The Pearson correlation coefficient is 0.40, which indicates weak correlation between the predicted LLPS propensity and concentration.

**Table 1.** The evaluation of the model proposed in this study on the test set. Data are presented as mean ± SD.

|  | Sensitivity | Specificity | Precision | Accuracy | F1 Score |
| --- | --- | --- | --- | --- | --- |
| Present model | 0.924±0.010 | 0.967±0.005 | 0.966±0.005 | 0.946±0.004 | 0.944±0.004 |

### LLPS propensity survey on the human proteome

A proteome wide LLPS propensity survey on the human proteome was performed, where a cutoff of 0.7 which was used by Hardenberg and colleagues for distinguishing LLPS driver and client proteins was firstly used. At this cutoff, ∼50% of the proteins are categorized as drivers. At a cutoff of 0.85, it dips to ∼35%. Both numbers agree with the aforementioned finding that LLPS is a proteome-wide phenomenon^13^. We are particularly interested in enrichment the high LLPS propensity (score > 0.85) and low propensity (score < 0.2) groups in any particular biological functions.

### Functional distribution of the high/low LLPS propensity proteins

The functional distribution of these two groups were performed by analyzing their GO terms (Fig. 1a, b). It is immediately evident that they are associated with distinguished GO terms. The group with a high LLPS propensity is dominantly enriched in mRNA processing and regulation related processes. While the other two terms are associated with the multicellular developmental processes. Enrichment of the high LLPS propensity proteins in the mRNA related processes is expected. There are rich types of nuclear condensate, e.g., PML bodies, nuclear stress bodies (nSBs) and transcriptional condensates. The enrichment in the developmental process (GO:0007389 pattern specification process; GO:0007351 regionalization) agrees with the observation that evolution toward higher organisms is associated with an increase of IDR regions and LLPS sequences. Some LLPS systems are known to be critical in the developmental processes, e.g., tight junction condensate^38^. For the high propensity group, the two most enriched terms are GO:0005667 (transcription regulator complex) and GO:0016607 (nuclear speck). For the low LLPS propensity group, the terms are notably enriched in a few amino acid and nucleotide related enzymatic functions. In terms of cell components, the core ribosome proteins and mitochondrial matrix proteins are known unlikely to be present in LLPS systems. Overall, the proteome-wide functional distribution of the groups cross-validated our LLPS propensity prediction algorithm. It also provides a glimpse of cellular protein partition and functions.

**Figure 1.**
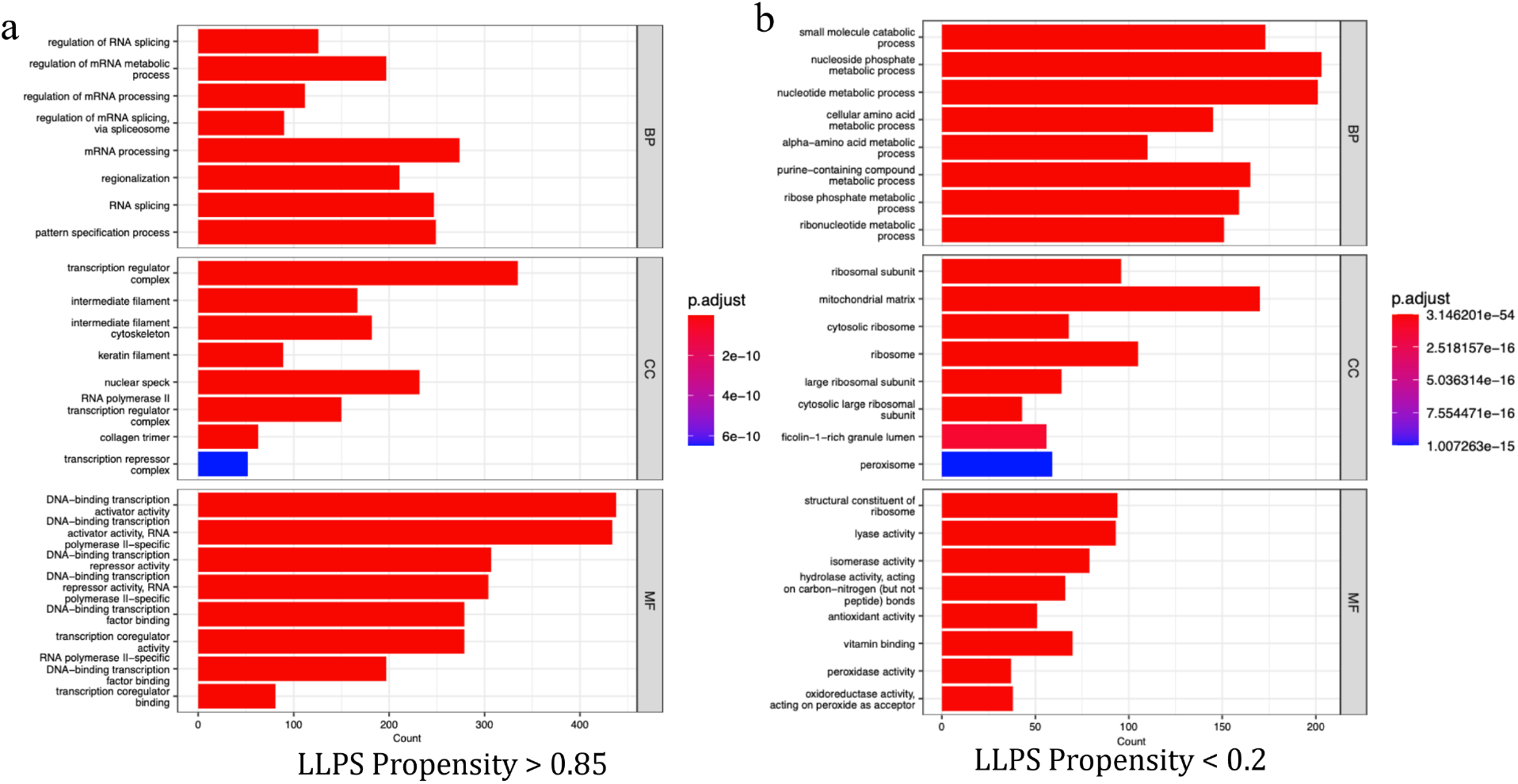
GO term enrichment analysis of human proteome. Bar graphs illustrating the Top 10 GO terms (MF = molecular function, BP = biological process, CC = cellular component) for: **a** high (score > 0.85) or **b** low (score < 0.2; right) LLPS propensity, respectively. P adjust value is a modified Fisher Exact P value for gene-enrich analysis.

### LLPS constituent prediction on specific systems

We firstly tested our LLPS constituent prediction algorithm on the SPOP-DAXX using speckle-type POZ protein (SPOP) as a query. The SPOP-DAXX is a nuclear condensate that can be formed under both *in vitro* and in cellular conditions^32^. The biological mechanism related with this system is that it serves as a hotspot in the nucleus for proteasomal labeling to the ubiquitination degradation. The SPOP-DAXX co-condensation mechanism has been studied *in vivo, in vitro*, and by biophysical techniques^32,39,40^. SPOP is considered as a tumor suppressor for regulating the degradation of numerous oncoproteins^4,5^. Its co-condensation with death-domain-associated protein (DAXX) triggers the polyubiquitination of DAXX and other target proteins. The presence of a long stretch of poly-glutamate sequence in DAXX may selectively attract DNA binding proteins, e.g., transcription factors, by its electrostatic property. It is proposed that, in the droplet state, the process involves the oligomerization of SPOP and the interaction between multivalent serine-rich (SR) linear motifs in DAXX and SPOP oligomer^39^. Certain cancer related SPOP variants deprive the protein of its LLPS capability, which is possibly the underlying pathological mechanism^32^. The SPOP-DAXX system requires additional proteinaceous components for the ubiquitination process as neither SPOP nor DAXX carries ubiquitin ligase activity. In this study, SPOP is the only input for the LLPS system reconstruction. In our algorithm, by the pairwise interaction between SPOP and DAXX, DAXX was identified as a LLPS driver protein and used as a query for the system expansion (Fig. 2a). From previous studies, it is known that SPOP is an adaptor protein for E3 ligase CUL3^41,42^. The addition of CUL3 is necessary for this system’s functionality, and our algorithm identified it in a tripartite interaction with SPOP as a bridge node. The SPOP-DAXX-CUL3 assembly features a combination of LLPS style interactions and specific PPI interactions, e.g., the SPOP-CUL3 interaction. Toward upper-right hand side of the network, the TP53-MDM2-USP7 module is vital for the homeostasis of cell proliferative state. MDM2 as a E3 ubiquitin ligase promotes the TP53 degradation, the deubiquitinase USP7 interacts with MDM2 to retard the auto-ubiquitination of MDM2, and DAXX-MDM2 co-localization enhances the E3 activity of MDM2 toward TP53^43^. The presence of this module in the LLPS system hints that it resembles the state of unstressed cells. The localization of both SPOP and DAXX to PML bodies has been well-documented^32^. Protein PML was seen as a top LLPS propensity node in the central part of the network. With PML, MDM2, and the left-hand side of PML, the organization of the network demonstrated the intriguing relationship between SUMO ligase activity and ubiquitin ligase activity. Among the eight nodes (RNF4, PIAS1, SIAH2, BRCA1, PML, MDM2, SUMO1, SUMO2), PIAS1 and PML carries SUMO ligase activity, while RNF4, SIAH2, BRCA1, MDM2 are RING-type E3 ligase. Both PIAS1 and PML were identified as LLPS drivers, together with DAXX, this system harbors a tripartite co-scaffold architecture. In addition to their transferase activity, PML, PIAS1, RNF4 as well as BRCA1 contains SUMO interaction motifs (SIMs)^44,45^. It has been proposed in previous studies, that the interaction between SUMO polymers and SIM-containing proteins is critical for the assembly and regulation of PML nuclear bodies^44,46^. Our predicted LLPS network is in line with this model, i.e., both PML and PIAS1 could spontaneously phase separate into the droplet. Their SUMO ligase activity can amplify the local valence concentration, i.e., SUMO concentration in this case, which in turn helps for the recruitment of other LLPS constituents, e.g., RNF4 and BRCA1. This cascade is previously referred to as the SUMO-targeted ubiquitin ligase (StUbL) pathway^47^. The presence of SIAH2-PIAS1 edge is possibly part of the regulation mechanism of this LLPS system. SIAH2 is proposed to mediate the degradation of PIAS1^48^. Moreover, the network is enriched with nucleotide binding proteins which concur with the nuclear location of the droplet. This network and LLPS constituent prediction feature a combination of LLPS-style and PPI-style interactions, which well illustrates both of the organization and regulation principles of the system.

**Figure 2.**
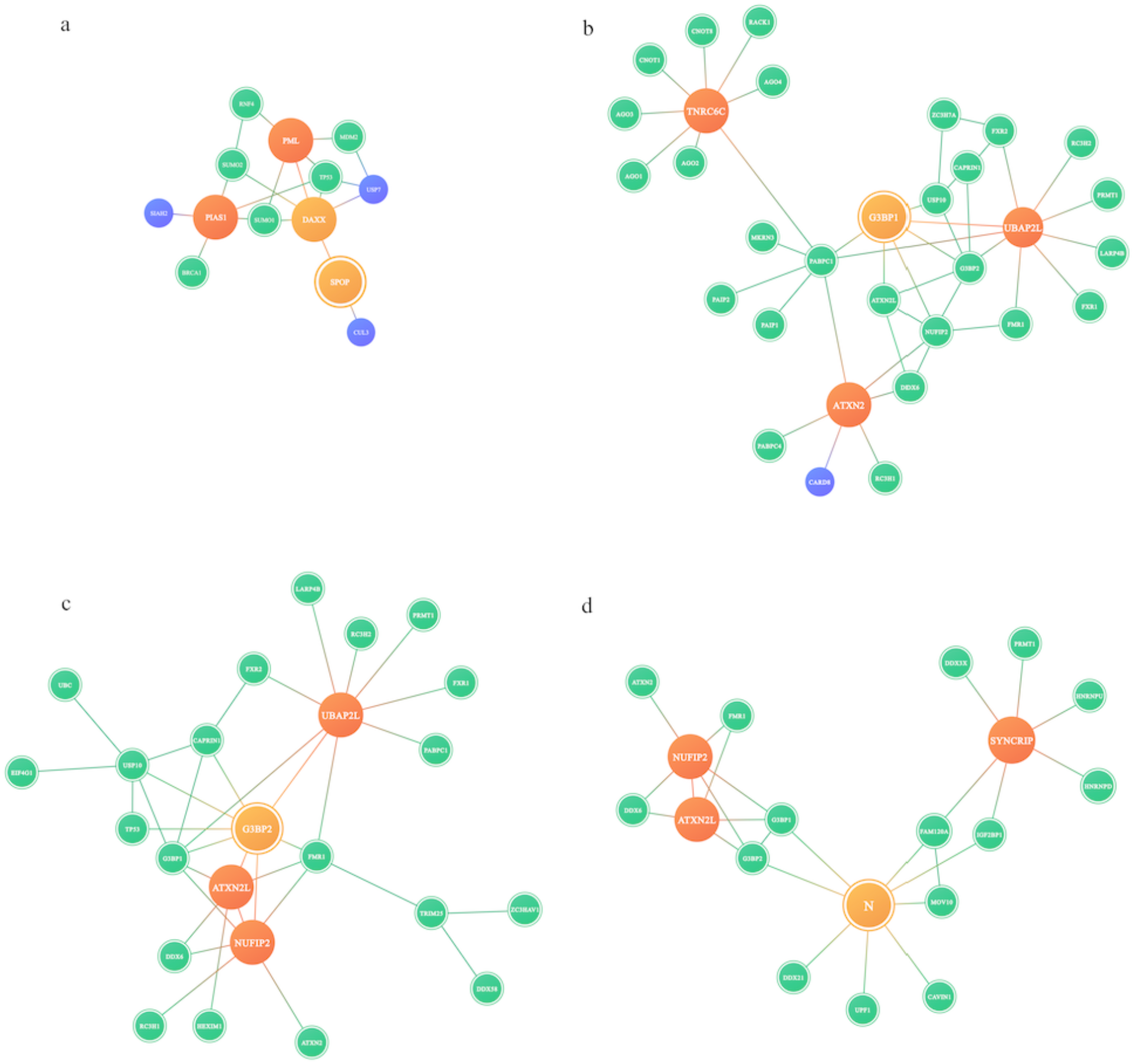
LLPS constituent prediction on specific systems. The LLPS system generated by our model. The blue circle represents a LLPS constituent with its name above it. The blue circle with green border represents constituent that is able to bind nucleic acid. The yellow circle represents proteins that are top LLPS scaffolds. The red circle represents other scaffold proteins. The yellow circle with red border represents the input query. **a** the SPOP-DAXX system with SPOP as initial query; **b** the SG system with G3BP1 as initial query, **c** G3BP2 as initial query; **d** SARS-CoV-2 nucleocapsid protein N LLPS prediction.

In the second and third case, we tested our algorithm on SG and SARS-Cov-2 protein N. The biological mechanism of SG is related with translational regulation and mRNA storage. In contrast to the SPOP-DAXX prediction, SG is known to be a comprehensive system that incorporates plenty of proteinaceous components and a significant percentage of the transcriptome. SG has been intensively studied, and its constituents are well documented^49-51^. On top of that, SG was used as a model system for studying organization principles of the droplet state, which leads to valence amplifier, bridge, and capping node notation of LLPS constituents^37^. Among the constituents, a few are principal and critical for the SG assembly, e.g., G3BP1, G3BP2, UBAP2L, etc. Both G3BP1 and G3BP2 are dimeric which is partially supported by their self-pairwise PPI record. Previous studies have shown that G3BP1 and G3BP2 double knockout cells are hampered in SG formation under As-induction^37^. We conducted two trials with G3BP1 and G3BP2 as query sequences, and the two networks are shown respectively in Fig. 2b, c. Although the two networks differ in their number of constituents (the G3BP1 network is more extensive), the two networks share very similar topology, especially for the eight constituents which partition strongly into G3BP NTF2 domain condensate^37^. They are G3BP1, G3BP2, USP10, UBAP2L, CAPRIN1, FMR1, FXR1, and NUFIP2. Additionally, ATXN2L, FXR2, DDX6, RC3H1, and RC3H2 are common between the two networks and are persistent constituents of SG. In the network constructed with G3BP1, two additional clusters of nodes that are respectively centered with PABPC1 and TNRC6C. Both proteins are known constituents of SG. TNRC6C is also known to be constituent of the P-body^52^. By comparing the two networks, core components of the network can be inferred, and it is in agreement with previous studies. In our design, we aim to keep the network concise and interpretable. The number of evaluated pairwise PPIs was thus limited to 10 to keep it as a relatively small number. This parameter can be readily adjusted to a larger number if a comprehensive network is more desirable. Although USP10’s presence is persistent and it is frequently involved in SG formation, its role is in controversy: in the work from Sanders et al.^37^, it is proposed as a capping node, which generally slows SG formation; Kedersha et al.^53^ propose that USP10/CAPRIN1 competes for G3BP binding and USP10 binding inhibits SG formation; Meyer et al.^54^ suggests that USP10 is critical for rescuing 40S subunits of ribosome from the SG. We observed that the USP10 region is a variable part of the network. Although both edges exist simultaneously between USP10/CAPRIN1 and G3BP, due to the ensemble nature of PPI data, it is not in conflict with the finding from Kedersha and colleagues. The presence of UBC and EIF4G1 as its neighbor hints that the functionality of this module is closely related with ubiquitination. This observation is independent from the later publication^54^ and demonstrates that our prediction indeed contains heuristic value within it. The DDX6 node is located closely to the key SG constituents, signifying that it is also closely related to SG regulation^55^, although its exact functionality is unknown.

The formation of SG is triggered by a variety of conditions including viral infection. From beginning of the 2019 SARS-CoV-2 outbreak, although lots of attention was paid to the spike protein, its virulence was instead closely related with its nucleocapsid protein N. N variants greatly enhanced the virus replication efficiency^36^. Under SARS-Cov-2 infection conditions, the SG formation is impaired^56^. This is proposed to be primarily mediated by protein N. We are interested in testing our algorithm under this varied condition. The BioGRID database makes a SARS-Cov-2 PPI dataset publicly available, and we queried protein N for prediction. The network prediction is shown in Fig. 2d. It is immediately evident that the network changed dramatically. Although part of the core SG network was unaffected (upper-left part), the UBAP2L cluster completely vanished. Toward the right-hand side, a bipartite interaction between N and G3BP1/G3BP2 replaced previous interactions. In the study from Sanders and colleagues, they found that the G3BP-UBAP2L interaction is critical for the SG condensation for UBAP2/2L 2KO leads to smaller SGs in a minority of cells^37^. Therefore, our observation follows the statement that protein N attenuates SG formation^56^. In the same study^56^, it also observed that G3BP1 is able to recruit more proteinaceous components than G3BP2, which is also noted in our predictions (Fig. 2b, c). In this combined case, we demonstrate that our algorithm can reliably identify critical components of the SG and when supplemented with additional data, it is able to probe system conversions.

In the fourth case, we applied our algorithm to the cyclic GMP-AMP synthase (cGAS) system. Protein cGAS senses the presence of poly-deoxynucleotides and forms droplets by multifaceted interactions with DNA^57^. Upon droplet formation, cGAS substrates, i.e., ATP and GTP are enriched into the condensate to produce cyclic GMP-AMP (cGAMP)^10^, which activates stimulator of interferon genes (STING) and in turn induces type 1 interferons. The cGAS-DNA condensation is proposed to be an efficient mechanism for cGAMP production. It is also known that cGAS could form condensate in both cytosol and nucleus. In the cytosol, cCAS can be activated by viral DNA or micronuclei^33^. Upon phosphorylation, cGAS is translocated into the nucleus^58^, where it exerts an enzymatic-independent function^59^. We tested our algorithm on this system for dissecting multi-location LLPS predictions. In the initial trial, a unified network was constructed (Fig. 3a) where the subcellular location was ambiguous and both nuclear/cytosolic constituents were permitted. Later, stringent subcellular location criterion was applied and led to two separate networks (Fig. 3b, c). The cytosolic network is minimal, harboring two LLPS scaffolds and another seven constituents. For the nuclear network, it contains significantly more members and three high LLPS propensity proteins. It is probable that upon translocation into the nucleus, cGAS and IFI16 can join some pre-existing nuclear condensates. cGAS-IFI16 interaction is persistent in both networks. In both networks, some key interactions are bridged by IFI16, e.g., cGAS-IFI16-STING and cGAS-IFI16-BRCA1 interactions. In the nuclear network, we identified RBBP8 as a constituent, i.e., an endonuclease which participates in DNA-end resection process. And LYAR is a negative regulator of innate immune responses. Although the nuclear cGAS function is still mystic, our prediction revealed some specific targets which facilitates its function by LLPS mechanism.

**Figure 3.**
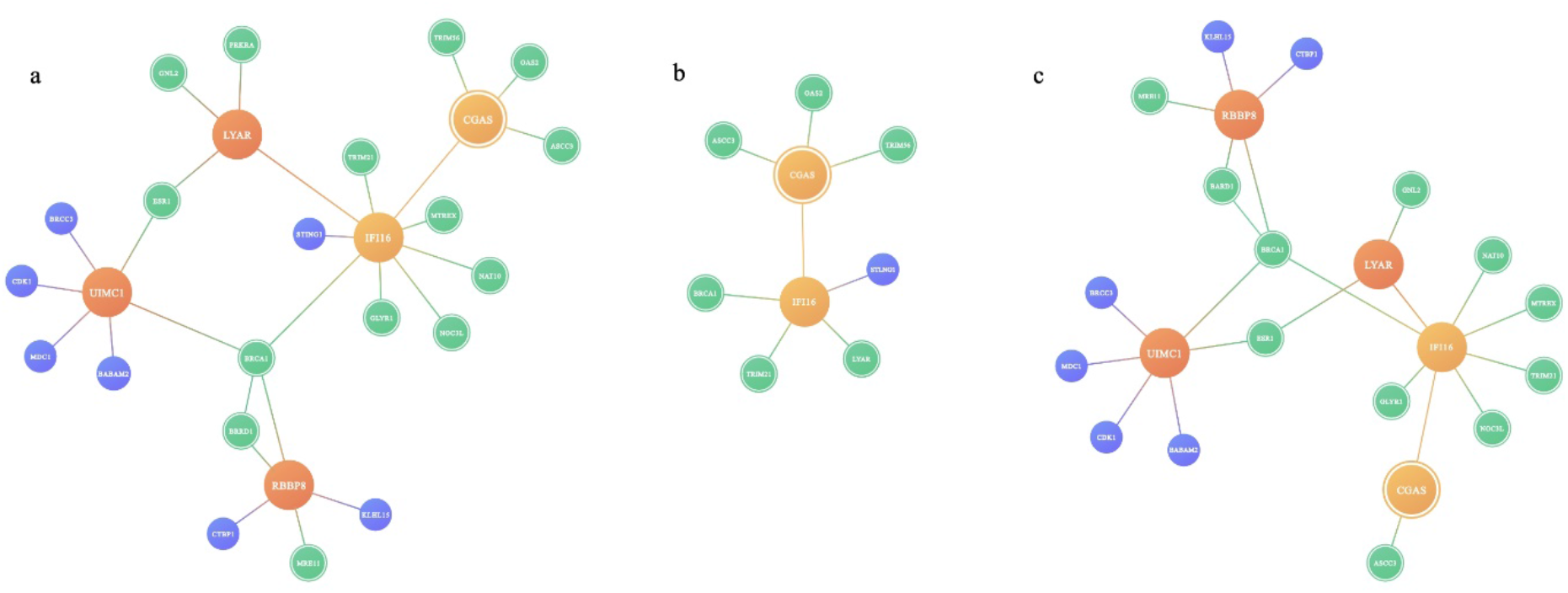
LLPS constituent prediction on the cGAS system. The cGAS LLPS system generated by our model. The color schema is the same as in figure 2. **a** the cGAS-STING system with cGAS as initial query, **b** the cytosolic network and **c** the nuclear network extracted from **a**.

For all of the four systems, detailed information of the identified constituents is provided in supplementary table 2-5.

## Discussion

We demonstrate that it is feasible to accurately predict the LLPS high frequency constituents by combining protein LLPS propensity prediction, PPI information, and spatiotemporal knowledge of the constituents with a few heuristic rules. The rules are inspired by our observations mentioned in the introduction and work from Sanders and colleagues^37^. In the work from Sander et al., the LLPS constituents are categorized into three types, i.e., valence amplifier (valence ≧ 3), bridge (valence = 2) and capping nodes (valence = 1). We hypothesize that for spontaneous LLPS proteins, their condensation increases the local valence concentration in a manner analogous to a self-amplifying chain reaction, and this set of proteins can be identified by LLPS propensity prediction. While being tricky to find a uniform propensity threshold that effectively differentiates LLPS drivers and non-drivers, with trial and adjustment, we demonstrate that a uniform threshold is feasible for our tested systems. The presence of bridge and capping nodes in SG is a reflection that LLPS interactions well embrace PPI style interactions, i.e., bridge and capping interactions that are essentially PPIs. This point of view is evident for proteins whose domains are feasible for structural studies, e.g., SPOP-CUL3 and other well-known adaptor domains^41^.

We assumed that there are a limited number of LLPS systems in the cells and their high frequency constituents are persistent. To impose this characteristic into our prediction, we built a bootstrap strategy into our algorithm. It is evident that a LLPS system could be co-scaffolded by more than one high LLPS propensity proteins, e.g., G3BP1, G3BP2 for the SG. This set of proteins are core components of the system. We limited the distance from any other nodes to any core components. In this way, a boundary condition is enforced. The default parameter for the maximum edge distance is three, which is envisaged with driver-bridge-bridge-capping, or driver-driver-bridge-capping style interactions. For top scaffold nodes, their direct PPIs are always kept. Although we do not have an explicit definition of bridge or capping nodes, the nodes toward the exterior of the network are of lower LLPS propensity by construction. Even when the very same set of rules and procedures are applied, the number of identified constituents are quite different for different systems. It essentially relates to the topology of the underlying PPI network, i.e., for some PPI network regions, e.g., SPOP-DAXX network, the PPIs are self-enclosed by mutual PPIs, while in some other regions, the PPI network is more extensive.

We also observed that for some systems there are driver-bridge/capping-driver style bipartite interactions. It can be rationalized with example as the SG and P-body have spatial adjacency relationship. In this study, we identify IFI16 which could locate the cGAS LLPS system toward pre-existing STING condensate. This form of bipartite interaction is therefore a feasible mechanism for LLPS system synergies. Some high propensity proteins, such as TP53, appear in many LLPS systems. The bipartite style LLPS interaction is present at high frequency when TP53 was involved. One possible rationale could be related with pathway cross-talks. As the goal of this algorithm is the construction of specific types of LLPS network, we cut the bipartite interaction at the far from query side to keep the constructed network concise and tractable for further analysis. In future studies, we could specifically investigate the relationship between different LLPS networks. Of note, LLPS systems bring up tangible scenarios where two types of interactions are highly relevant, i.e., transient interactions and indirect interactions. Transient interactions are highly relevant in protein post-translational modifications and signaling processes. Studies on LLPS systems show that proteins can be brought into spatial proximity even if their interaction is not mutually coded by their sequences. The enrichment of these two types of interactions in the droplet state demonstrate how emergent properties are fostered by the equipment of LLPS. It is also bearing in our mind that LLPS regulation mechanisms are feasible for the regulation of these two types of interactions.

Another inspiration of this study came from the hypothesis of LLPS-PPI co-evolution. By analyzing the amino acid frequencies in the eukaryotic IDR region^26^, they are surprisingly similar with LLPS sequences^8^. Given that the arise of eukaryotic organism is an early event in evolution (estimated to be at least 2.7 billion years ago^60^), if the equipment of LLPS was harvested by the cell not much later than the event, LLPS systems and PPI networks would have long history of coordinated coevolution^28^. For the capability of LLPS on the spatiotemporal regulation of protein distribution and its proteome-wide spreading, PPI and LLPS networks are deeply engaged and have an intimate relationship. Our findings in the study lend support to this idea.

Using aggregated PPI data, i.e., the PPI terms in most cases are compiled from more than one condition, for practicality as evaluating the confidence of a specific PPI requires a sufficient amount of data, our method can reliably identify high frequency LLPS constituents. It should be mentioned that for the SG constituents’ prediction, the conditions are not strictly relevant for the SG formation; however, the PPI network is still expedient for the algorithm. Possible explanations are that some LLPS systems are seeded before induction or LLPS interactions might be formed during the AP-MS process. In future studies, it would be interesting to combine computational methods, such as the method presented in this study, and PPIs that are measured from specific conditions, e.g., specific cancer cell lines instead of from workhorse cell lines^29-31^, to see whether PPI network changes could be related to LLPS system changes.

The cellular environment has long been known to be crowded with different types of biomolecules^61^, and LLPS systems bring a novel sight to this ‘functional crowding’. While crowding is a generic feature of LLPS systems, physiological LLPS systems can cluster a specific set of constituents. In this study, we present a first-in-class approach of LLPS constituents’ prediction and have demonstrated the effectiveness of this approach. Given desires for a tool that can help on the rational design of LLPS related experiments, our method that can be readily extended to the proteome scale gains even more value, since in creating a picture of the cellular partition landscape, we will have taken a large step forward in the study of the cell^1^.

## Methods

### Preparation and construction of datasets

Two types of datasets have been explored in this study: a dataset for training the LLPS propensity predictor and an aggregated experimental PPI dataset. The sequence and sequence annotation, including subcellular location and function information on the target proteins, is fetched from the UniProt database^62^ in an *ad hoc* manner through web service.

The LLPS propensity predictor training dataset is made up of three subsets. The training set is used for the training and validation of the propensity predictor, which is adapted from Saar et al.^16^. In this study, the former LLPS+ and LLPS-set is merged into a single set. The former +/-set was divided on the setting of a threshold concentration of 100 *μ*M, however, we found the division was difficult to relate with the *in vivo* conditions.

The PDB* set was essentially unchanged. The other two subsets were used for the benchmark of the prediction algorithm: the benchmark set used by the DeePhase method; the A-IDP set which is a novel test dataset that we compiled from Dzuricky et al.^8^. In the study, the upper critical solution temperature (UCST) of a series of synthetic LLPS sequences was probed systematically. The sequences are homotypical and contain no-posttranslational modification, and therefore are suitable for benchmark.

In this study, we relied on the latest release of BioGRID^63^ for the PPI dataset. The BioGRID raw data was downloaded in *tab3* format and the individual pairwise interactions were aggregated. The processed BioGRID data was used as input for LLPS constituent prediction algorithm.

### LLPS propensity prediction algorithm

The deep learning (DL) approach is taken to predict the phase separation propensity of the target protein. The DL model takes one protein sequence as input and predicts its likelihood of spontaneous demixing. After merging the former LLPS+ and LLPS-set, it was applied as the positive dataset for the training and validation, and PDB* was applied as the negative dataset. Protein sequence embeddings were evaluated using a pre-trained ESM model^64^. The ESM approach explored datasets with up to 250 million protein sequences using the masked language modeling and has been previously shown to capture biochemical properties of amino acids such as hydrophobic, polar, and aromatic groups etc. This preprocessing method generated fixed-length per-sequence representations. 1280-dimensional embedding vectors were adopted from variable-length sequences using a pre-trained ESM-1 model, which can be natively taken as input into the DL model. A fully connected neural network was implemented in PyTorch and trained with default parameters. The hyperparameter tuning was performed with a little improvement in accuracy. It has been trained for 200 epochs with the Adam optimizer, a 32-batch size, and a learning rate of 0.005. The training dataset was randomly split into a train and a validation set in a 1:9 ratio and the validation set was used to estimate the performance of the model, which was evaluated based on specificity, sensitivity, accuracy, and F1-score. The final prediction results were compared with the DeePhase approach on both validation and test sets^16^.

### Human GO term enrichment analysis for LLPS proteome

GO analyses were performed to capture the cellular component (CC), molecular function (MF), and biological process (BP) terms for picturing the functional distribution of LLPS proteome (LLPS score > 0.85) and non-LLPS proteome (LLPS score < 0.2). Gene set enrichment analysis (GSEA) was applied to annotate the enriched GO modules with Fisher’s exact test *p*-value < 0.05 for both groups using the “ClusterProfiler” R package^65^ to find which specific biological annotations are most closely related to different set. GO terms were ranked according to their enrichment scores.

### LLPS constituent prediction algorithm

In our algorithm (Fig. 4), the user can initiate the query with any protein sequence or UniProt ID. The first step in the protocol is to calculate the LLPS propensity of the query protein. If the query sequence’s LLPS propensity is lower than the ‘driver’ threshold (LLPS score < 0.85), the algorithm will search its direct PPI partners for any LLPS driver proteins. If there are none, the search will end and conclude that the query protein is not a constituent of an LLPS system. If direct PPI partners of the query contain one or more driver proteins, the one with the highest LLPS propensity score will be used as the query sequence. This new query sequence will identify its high frequency direct PPI partners. If the initial query has a propensity higher than the threshold, the algorithm will directly start with its PPI partner search. The number of occurrences of a pairwise PPI is read, in a loose sense, as confidence of the interaction. For the absolute majority of the proteins, their known PPI partners are on the order of hundreds. By default, we limited query protein’s number of PPI partners to 10, but it is kept as a variable that the users can explore. After PPI partners are identified, the sequence annotations filter out PPIs that are hardly physiologically relevant, e.g., cellular sublocation mismatch or molecular function/biological process mismatch. Notably, a protein could be annotated with multiple subcellular locations. In this case, as long as one of the locations matches, it is deemed that PPI is feasible.

**Figure 4.**
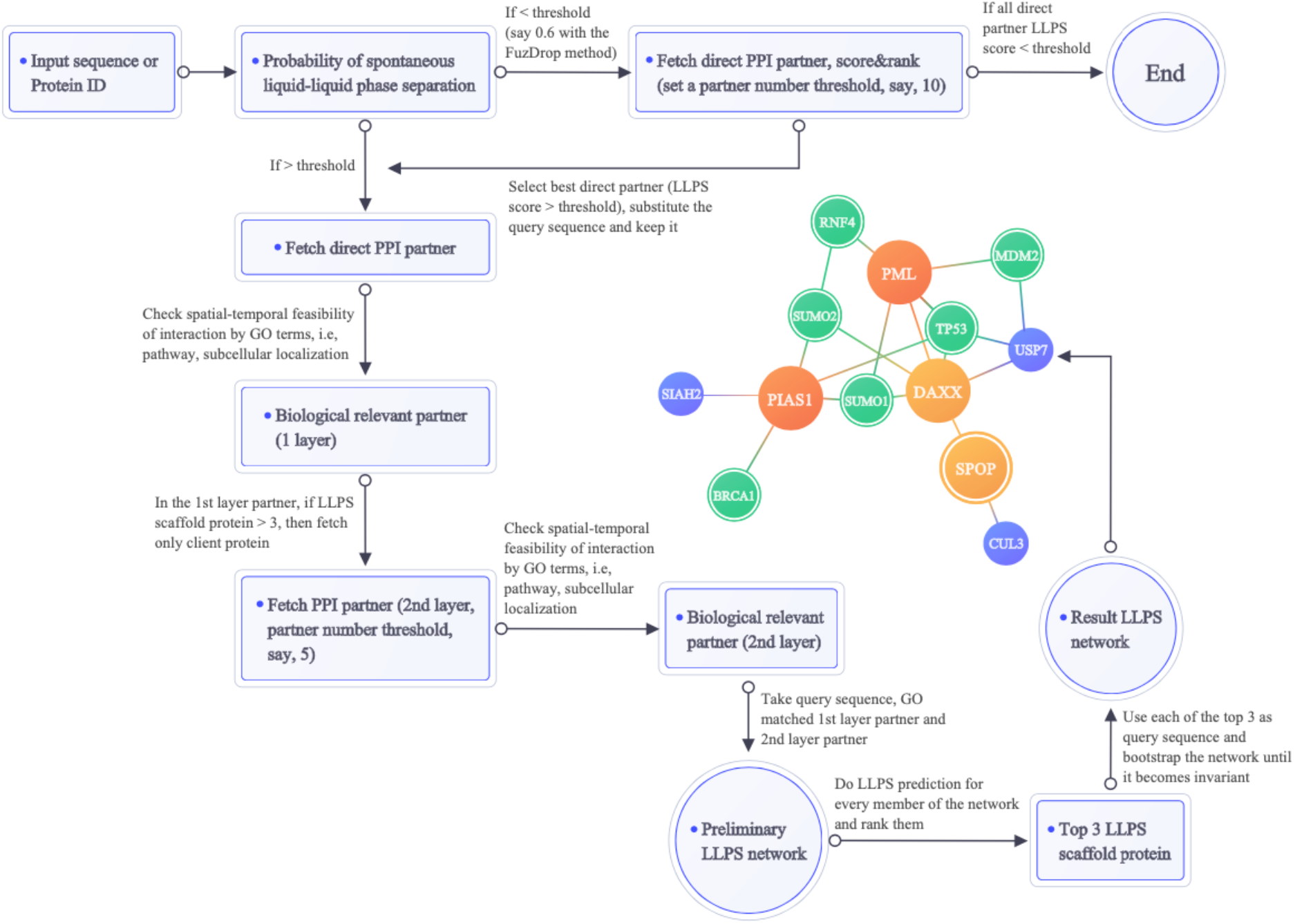
The LLPS constituent prediction workflow.

With the list of filtered PPI partners, two operations are carried out: 1) LLPS propensity calculation on each target, and 2) a PPI partner search with each partner as a new query. For second-layer PPI partner search, the hit number is limited to five, and for each hit, the annotation-based filtering procedure was applied. When a preliminary LLPS constituent list is assembled, the top three hits with the highest LLPS propensity can be identified. It is followed with a procedure which we termed “LLPS system bootstrap”. In this procedure, each of the top three hits was used as the query sequence, the subsequent three networks were combined. The unified network is trimmed by constraining the edge distance from any of the top hits or the initial query. This procedure reduces the dependency on the initial query sequence. In the method setup run, we trailed with two criteria for the second-layer hits, i.e., whether the high propensity hits are kept or discarded. We found that by keeping the second-layer high LLPS propensity hits, we could acquire more information on the relationship between different LLPS networks. Thus such hits are kept in the current algorithm.

## Data Availability

All data within this paper are available from: https://github.com/promethiume/LLPSWise

## Code Availability

All codes are available from: https://github.com/promethiume/LLPSWise

## Acknowledgements

We thank J. Zhou, X. Xiang, S. Guo, G. Peng, Y. Xin, L. Wei, T. Tao, H. Zheng, Y. Yang, X. Li for their contributions. Y. Yang, X. Li and the Product Development team for help with the figures; G. Peng, Y. Xin, L. Wei, T. Tao, H. Zheng and the Innovation Center for their helpful discussions and support; J. Zhou, X. Xiang, S. Guo and our colleagues at StoneWise AI for their vision and encouragement.

## Supplementary Information

Supplementary Information: should be combined and supplied as a separate file, preferably in PDF format.

